# Intercalated Disk Nanoscale Structure Regulates Cardiac Conduction

**DOI:** 10.1101/2021.04.13.439660

**Authors:** Nicolae Moise, Heather L. Struckman, Celine Dagher, Rengasayee Veeraraghavan, Seth H. Weinberg

## Abstract

The intercalated disk (ID) is a specialized subcellular region that provides electrical and mechanical connections between myocytes in the heart. The ID has a clearly defined passive role in cardiac tissue, transmitting mechanical forces and electrical currents between cells. Recent studies have shown that Na^+^ channels, the primary current responsible for cardiac excitation, are preferentially localized at the ID, particularly within nanodomains around mechanical and gap junctions, and that perturbations of ID structure alter cardiac conduction. This suggests that the ID may play an important, active role in regulating conduction. However, the structure of the ID and intercellular cleft are not well characterized, and to date, no models have incorporated the influence of ID structure on conduction in cardiac tissue. In this study, we developed an approach to generate realistic finite element model (FEM) meshes replicating ID nanoscale structure, based on experimental measurements from transmission electron microscopy (TEM) images. We then integrated measurements of the intercellular cleft electrical conductivity, derived from the FEM meshes, into a novel cardiac tissue model formulation. FEM-based calculations predict that the distribution of cleft conductances are sensitive to regional changes in ID structure, specifically the intermembrane separation and gap junction distribution. Tissue-scale simulations demonstrated that ID structural heterogeneity leads to significant spatial variation in electrical polarization within the intercellular cleft. Importantly, we find that this heterogeneous cleft polarization regulates conduction by desynchronizing the activation of post-junctional Na^+^ currents. Additionally, these heterogeneities lead to a weaker dependence of conduction velocity on gap junctional coupling, compared with prior modeling formulations that neglect or simplify ID structure. Further, we find that disruption of local ID nanodomains can lead to either conduction slowing or enhancing, depending on gap junctional coupling strength. Overall, our study demonstrates that ID nanoscale structure can play a significant role in regulating cardiac conduction.

## Introduction

The intercalated disk (ID) is the specialized structure at the site of the cell-cell interface connecting myocytes in the heart, facilitating electrical coupling via gap junctions (GJs) and mechanical coupling via mechanical junctions (MJs) [1]. Recent evidence has high-lighted the structural heterogeneity of the ID [2, 3, 4, 1, 5, 6, 7, 8]: The ID consists of highly tortuous plicate regions, which are oriented perpendicular to the myocyte’s axis, are comprised of membrane “folds,” and contain the majority of MJs (adherens junctions, desmosomes, and composite junctions); and interplicate regions, which run parallel to the myocyte’s axis and contain most GJs. MJs and GJs have clearly defined roles in the pas-sive conduction of mechanical forces and electrical currents, respectively. However, recent ιι studies show that multiple electrogenic proteins regulating conduction, in particular the voltage-gated Na^+^ channel, Na_V_1.5, are localized at the ID [9, 10, 11,12,13,14, 15, 16, 17], in close proximity to the junctions, specifically GJ-adjacent perinexi and MJ adhesion-excitability nodes, forming specialized nanodomains in the intercellular cleft spaces [18]. Further, structural perturbations of the ID have been shown to alter conduction and risk of arrhythmias [19, 20, 21, 22, 10], and ID structure is disrupted in patients with atrial fibrillation [23]. Collectively, these findings suggest that ID structure plays an important active role in regulating cardiac conduction.

Standard cardiac tissue modeling approaches, e.g., the monodomain model, typically neglect ID structure and non-uniform ion channel distributions. However, prior *in silico* studies, including our work, that do account for these details predict that Na_V_1.5 localization at the ID facilitates cell-cell coupling via a mechanism known as ephaptic coupling [24, 25, 26, 27, 28, 29, 30, 31, 32, 33, 34, 35, 9, 36]. Ephaptic coupling is mediated by interactions between Na^+^ currents, *I_Na_*, on adjacent ID membranes sharing a restricted intercellular cleft space and the associated electrochemical gradients that form in this space. Specifically, during the cardiac action potential, I_Na_ at the ID in a depolarizing cell (i.e., pre-junctional *I_Na_*) reduces the electrical potential of the intercellular cleft. This reduction in intercellular cleft potential depolarizes the apposing cell from the extracellular, rather than the intracellular, side of the cell membrane. Additionally, ID-localized *I_Na_* reduces the local Na^+^ concentration of the intercellular cleft, and collectively these electrochemical gradients alter both the apposing ID membrane transmembrane potential and post-junctional *I_Na_* driving force. Prior work has predicted that a narrow intercellular cleft width, which enhances the effects of ephaptic coupling, can either enhance or slow conduction, depending on the timing and magnitude of post-junctional *I_Na_* [27, 25, 32].

While providing important insights into how non-uniform Na^+^ distribution regulates conduction, most of these modeling studies simply represent the intercellular cleft as a single, uniform compartment. Two recent studies have considered spatial variation within the ID and the cleft: Mori et al [25] investigated ephaptic coupling in a radially-symmetric strand of cells, while Hichri et al [37] modeled the ID Na^+^ channel distribution as either uniform or in two apposing clusters. However, all of these prior studies make a key simplifying assumption about the ID and intercellular cleft space, specifically that the cleft is a uniform, cylindrical space and the intermembrane separation between apposing ID membranes is uniform throughout the ID. Thus, all prior modeling studies have neglected the structural heterogeneity of the ID and intercellular cleft space (e.g., variations between plicate and interplicate regions). Thus, the mechanisms underlying conduction changes due to ID structural perturbations [19, 20, 21, 22, 10] are not well understood.

The goal of this work is to develop an approach to integrate ID structural heterogeneity into a model of cardiac tissue and investigate the impact of this heterogeneity on conduction. For this approach, we first generate realistic 3D finite element model (FEM) meshes of the ID and intercellular cleft, based on measurements from transmission electron microscopy (TEM) images of mouse ventricles. We then calculate a reduced electrical network, based on the FEM meshes, to represent the cleft, which crucially bypasses the computationally costly step of simulating electrostatics on the full 3D FEM mesh, and incorporate this reduced network into a novel cardiac tissue model formulation that accounts for the heterogeneity at the ID. In our study, simulations predict that ID structural heterogeneity has a significant impact on electrical conduction. Notably, we find that both the monodomain and single cleft tissue models either over- or underestimate predictions of conduction velocity (CV), depending on the relative strength of GJ coupling. Importantly, the ID structure results in heterogeneous, asynchronous behavior of the electrical potential and Na^+^ concentration in the intercellular cleft and *I_Na_* at the ID, and these changes collectively regulate cardiac conduction. These findings improve our understanding of arrhythmia mechanisms associated with pathological ID structural remodeling and suggest the ID structure as a potential new therapeutic target.

## Methods

The overall goal of incorporating ID structure into a cardiac tissue model is accomplished in three stages. First, we develop an algorithm to construct a 3D finite element model (FEM) mesh of the ID and intercellular cleft, reproducing the structure of key ID measurements from transmission electron microscopy (TEM) images. Importantly, this mesh generation incorporates several orders of magnitude of structural details, from the nanoscale structure of intermembrane separation, up to the microscale structure of the interplicate and plicate regions. Second, we calculate an equivalent electrical network for the conductivity within and out of the intercellular cleft space. We calculate this reduced cleft network by partitioning the full finite element mesh into a tractable number (25-200) of compartments and determining the equivalent electrical conductance between all pairs of adjacent compartments. Finally, we incorporate this cleft network into a cardiac tissue model, in which neighboring myocytes are coupled via GJs and their shared intercellular cleft space. Importantly, by representing ID-localized ion channels that induce heterogeneous electrical polarization and ionic concentration gradients within the intercellular cleft spaces, the tissue model links nanoscale ID structure with macroscale cardiac tissue function.

### Tissue Collection

All animal procedures were approved by Institutional Animal Care and Use Committee at The Ohio State University and performed in accordance with the Guide for the Care and Use of Laboratory Animals published by the U.S. National Institutes of Health (NIH Publication No. 85-23, revised 2011).

Male C57/BL6 mice (30 grams, 6-18 weeks) were anesthetized with 5% isoflurane mixed with 100% oxygen (1 l/min). After loss of consciousness, anesthesia was maintained with 3-5% isoflurane mixed with 100% oxygen (1 l/min). Once the animal was stably in a surgical plane of anesthesia, the heart was excised, leading to euthanasia by exsanguination. The isolated hearts were prepared for fixation for Transmission Electron Microscopy (TEM). Ventricles were dissected and fixed overnight in 2% glutaraldehyde at 4°C for resin em-bedding and ultramicrotomy as previously described [20, 10].

### Transmission electron microscopy and quantification

Transmission electron microscopy (TEM) images of the ID, particularly gap junctions (GJs) and mechanical junctions (MJs), were obtained at 6,000x, 10,000x, and 20,000x magnification on a FEI Technai G2 Spirit transmission electron microscope (ThermoFisher Scientific, Hillsboro, OR). Morphometric quantification was performed using ImageJ (NIH, http://rsbweb.nih.gov/ij/) by manually identifying and quantifying the following 11 specific ID measurements. Images at 6,000x magnification were used to quantify: (i) total ID cross-sectional length. Images at 10,000x magnification were used to quantify: (ii, iii) the length of plicate and interplicate regions (Fig 1B); (iv, v) the amplitude and frequency of plicate folds (Fig 1E), and (vi-ix) the lengths of GJs and the fraction of membrane comprised of GJs, in both plicate and interplicate regions (Fig 1E). Images at 20,000 magnification were used to quantify: (x, xi) the inter-membrane distance in the plicate and interplicate regions, identified by regions outside MJs and GJs, respectively (Fig 1E). The Wilcoxon rank sum test was used for single comparisons between plicate and interplicate region measurements. A *p* < 0.01 was considered statistically significant. Measurements are reported by the mean ± standard error, and with the first and third quartile range.

**Figure 1.**
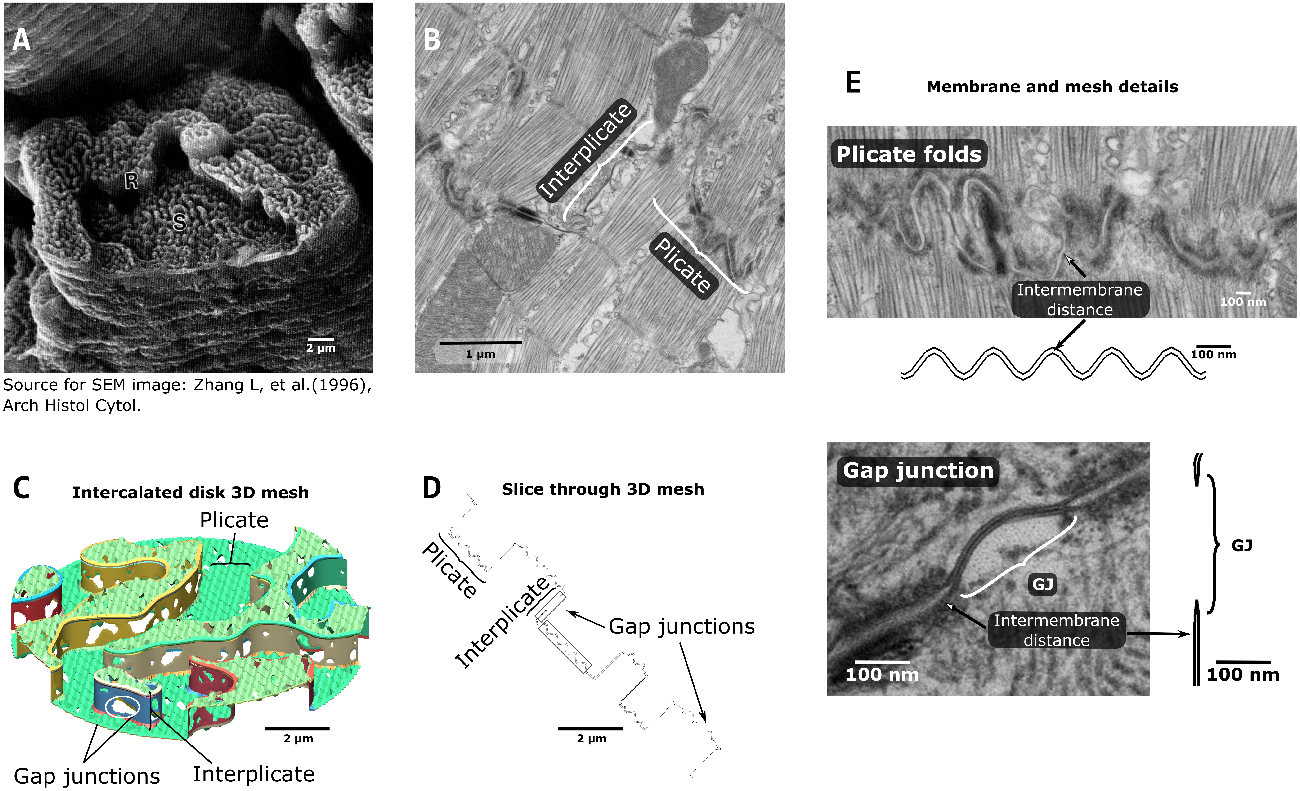
Intercalated disk structure and finite element model representation. **A**. Scanning electron microscopy (SEM) image of the intercalated disk from monkey ventricular myocyte. S and R indicate ‘steps’ and ‘risers,’ denoting the plicate and interplicate regions, respectively. Reproduced from [6] with permission. **B**. Transmission electron microscopy (TEM) image of mouse ventricular myocyte ID, illustrating the plicate and the interplicate regions. **C**. View of the 3D finite element model (FEM) model, with plicate and interplicate regions and gap junctions (GJs) highlighted. **D**. Slice through the FEM mesh, illustrating a corresponding view to the 2D TEM image. The sections in the grey boxes are enlarged in panel E. **E**. Magnified view of TEM images and FEM mesh slices, showing structural details for the plicate folds (top) and the gap junctions (bottom).

### Finite element model mesh generation

Based on measurements from TEM images, we developed an approach to replicate the nanoscale structure of the ID, generating realistic 3D finite element model meshes of the two cell membranes at the ID and the enclosed intercellular cleft space (Fig 1). Ŭe resulting structures are randomly generated but parametrically defined based on the TEM measurements described above (Fig S1, left). Specifically, these mesh parameters define the key features of the ID structure and fit them to experimental data: ID diameter, plicate and interplicate region length, plicate and interplicate region intermembrane distance, GJ size and distribution in both regions, and plicate fold amplitude and frequency (Table S1).

### Intercalated disk and gap junction map generation

The first step in the mesh generation involves developing “maps” that represent the patterns for two key structures, specifically the plicate regions within the ID and location of GJ clusters (Fig 2). Full details of this map generation approach are described in Supporting Material. The ID map represents a cross section of the entire ID, such that the map defines different “levels” or “tiers” of plicate regions (defined as representing the (x, y) plane), which are separated by interplicate regions (in the *z*-direction) (see Fig 1C). The GJ map represents the locations of individual GJ clusters. The same methodology is used to generate maps of both structures, with different algorithm parameters. In brief, a 2D map is generated by filtering and thresholding Gaussian random noise (Fig S2A), for which the threshold defines two sets of clusters (plicate levels for the ID map, GJ vs membrane for the GJ map). By varying the properties of the Gaussian noise and filter, we vary the properties of the resulting maps (Fig. S2B, C). Importantly, we adjust mesh generation parameters (Table S1) such that the map properties match the corresponding TEM measurements, specifically measurements of plicate region length, and GJ lengths and distributions in the plicate and interplicate regions. In particular, estimating the map generation parameters required comparing the 2D map properties with the 1D measurements obtained from TEM images; we performed “slices” of the 2D maps to obtain a series of 1D length measurements along the slice to directly compare model and experimental measurements (illustrated in Figure S3). Additionally, the random nature of the algorithm is a key feature of the mesh generation and overall study, as this enables the generation of multiple IDs with comparable and statistically similar overall properties and allows us to account for experimental variability in ID structure.

**Figure 2.**
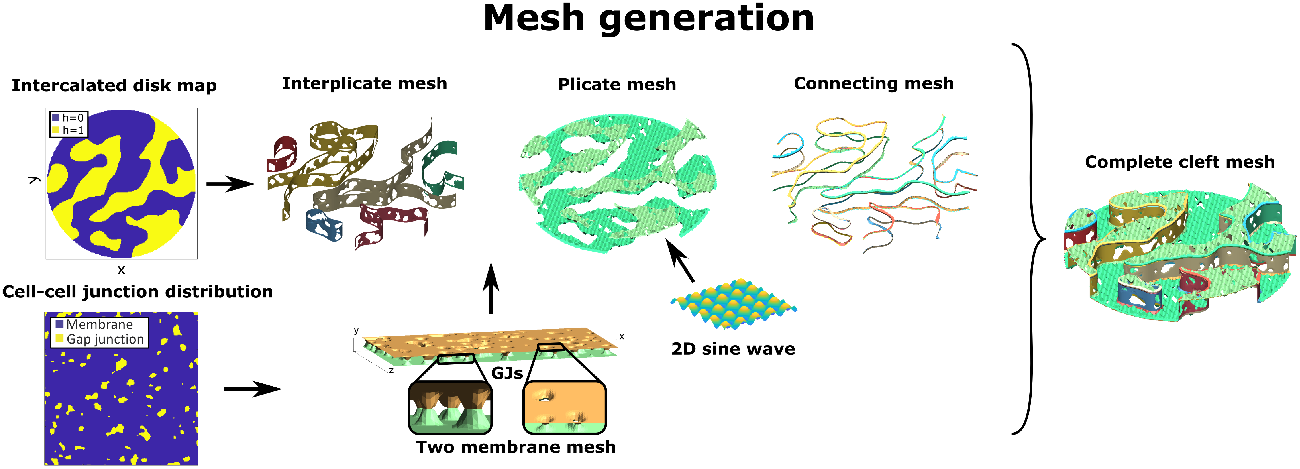
Outline of finite element model mesh generation algorithm: The intercalated disk (ID) map defines two “levels” of plicate regions, such that the interplicate region mesh is defined by the region separating the two levels. The cell-cell junction distribution determines a GJ map for both plicate and interplicate regions, which defines the locations of the two membranes connecting. A 2D sine wave function replicates the folds in the plicate membranes. A series of connecting meshes join the plicate and the interplicate region membranes, ensuring that the full mesh encloses a continuous volume (i.e., the mesh is “waterproof”). Combining all of these mesh components generates the cleft mesh.

### Finite element mesh construction

After generating maps of the individual ID structures fit to TEM measurements, we next generate the finite element mesh, which is comprised of plicate regions, interplicate regions, and small connections between these regions (Fig 2). Each plicate and interplicate mesh is initially defined by two rectangular surfaces, which represent the two apposing cell membranes, and then is “populated” by GJs, with distribution specific to that ID region and determined independently. As GJs mediate direct electrical communication between cells, at the location of GJs, the two apposing cell membranes connect, which appear as “holes” in the mesh (Fig 1C-E). Additionally, the intermembrane distance between the two membranes is defined, based on TEM measurements, denoted as *d_P_* for the plicate region and *d_I_p__* for the interplicate region. Importantly, we note that these two distances can be independently varied.

We next transform the individual mesh components into the shape of the ID: The plicate meshes are cropped into the shapes of the individual levels, determined by the ID map, and further are also “folded” using the shape of a 2D sine wave, to match the amplitude and frequency of the plicate folds (Fig 1C-E). The interplicate region meshes are rectangles, with lengths that match the contours of the plicate region levels and height determined by the TEM measurements of interplicate region length. The interplicate meshes are then curved and rotated to be positioned between the corresponding plicate region meshes. Small mesh pieces then join all adjacent plicate and interplicate regions. This complete mesh represents the two apposing cell membranes. The mesh is then checked and repaired using the meshfix algorithm [38], ensuring that the surfaces completely enclose a volume, representing the intercellular cleft. Finally, we use Gmsh, a finite element mesh generation software [39], to generate a 3D tetrahedral mesh of the intercellular cleft space.

### Intercellular cleft conductance measurement

We next developed an approach to reduce the full 3D finite element mesh of the intercellular cleft space to an equivalent electrical network (Fig S1, right). Network connections represent the electrical conductivity between adjacent regions in the intercellular cleft and are calculated by solving the full electrostatics problem on each pair of adjacent cleft compartments. This simplification facilitates incorporating the reduced cleft network into a cardiac tissue model, thus integrating the nanoscale detail of the ID into a tissue-scale model (Fig 3). The finite element mesh is first divided in *M* compartments, where *M* is a tractable number between 25-200, using the Metis partitioning algorithm [40] in Gmsh, which generates partitions of approximately equal volume and well-behaved boundaries (Fig 3A). We next calculate the centroid of each compartment, which represent the location of the cleft network nodes (Fig 3D, G (right)). For each pair of adjacent compartments (*j* and *k*), we “cut” (dashed lines) and generate a new finite element mesh of the shared volume between the two centroids (Fig 3B). To calculate an equivalent conductance between the adjacent nodes, we solve the full 3D electrostatics problem on this new mesh using the finite element solver in MATLAB (The Mathworks, Natick, MA); specifically we solve Laplace’s equation for electrical potential (Fig 3C), given by the following partial differential equation:

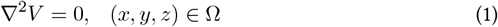

where *V*(*x*,*y*,*z*) is the electrical potential on the shared volume mesh, and Ω is the domain of the shared volume. The boundary conditions are defined such that *V* = *V_fem_* on one of the cut surfaces running through a centroid (dashed line in Fig 3B), and *V* = 0 on the other cut surface. Using the numerically-calculated potential V, we next calculate the total current between the two boundaries through the mesh, *I_fem_*, given by the following surface integral over one of the cut boundaries,

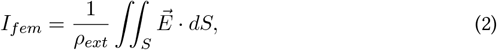

where 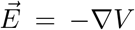 is the electrical field, *ρ_ext_* is the resistivity of the extracellular cleft space, and *S* is one of the cut surfaces on the boundary. Numerically, we calculate this integral by summing over all *N* triangular finite element faces on the boundary *S*,

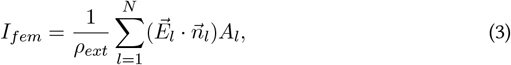

where 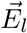 is the electrical potential gradient, averaged over the 3 vertices of the *l*th element face, 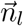 is the unit vector normal or perpendicular to *l*th element face, and *A_l_* is the *l*th element face area. We note that since Laplace’s equation is linear, the conductance value calculations do not depend on either which boundary surface is used for the current calculation, or the value of *V_fem_*. Thus, for simplicity, we set *V_fem_* = 1 V.

**Figure 3.**
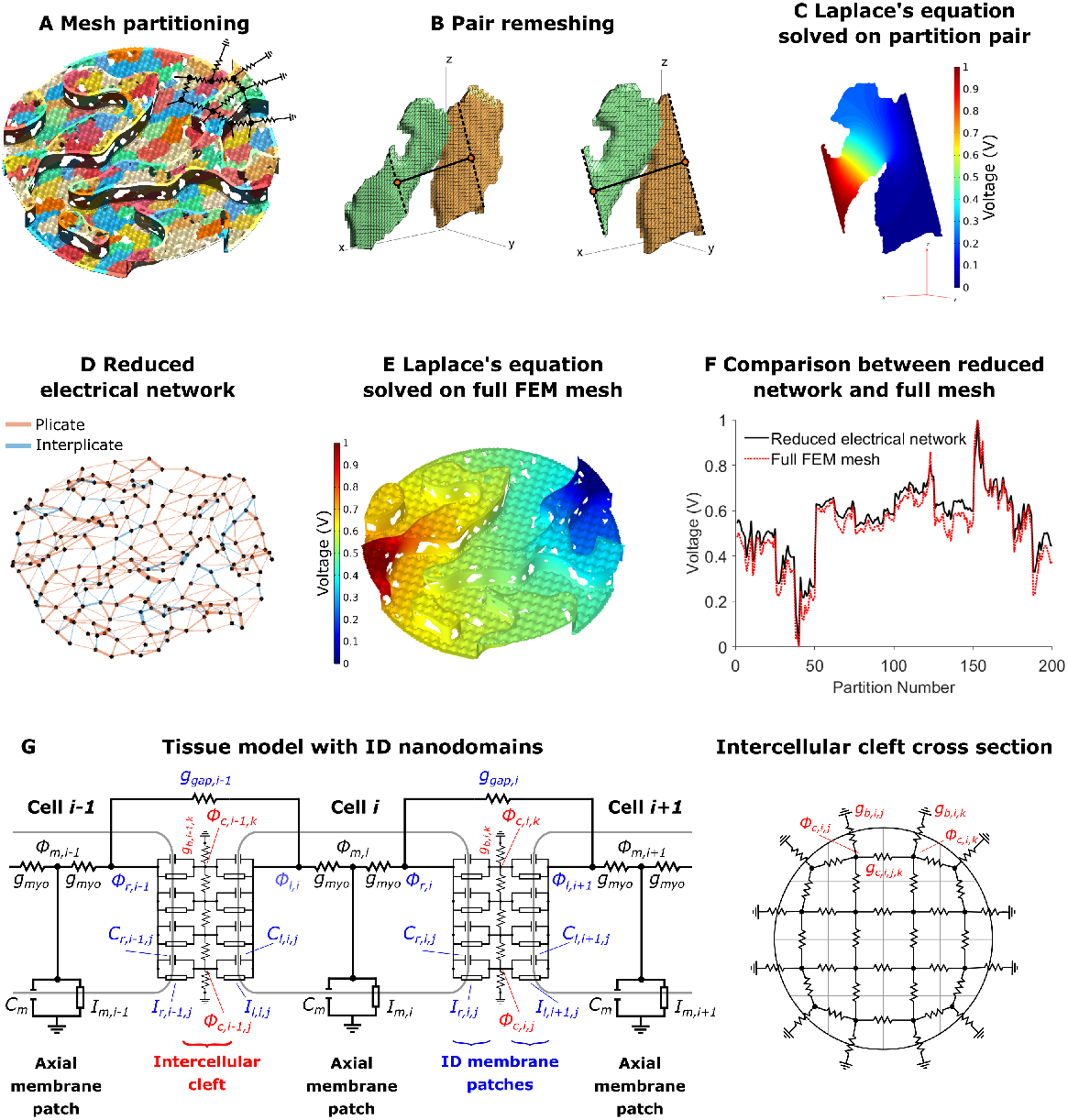
Intercellular cleft conductivity and tissue model with intercalated disk (ID) nanodomains. **A**. Partitioning of the finite element model (FEM) mesh into 200 partitions or compartments, with different colors denoting different compartments. Overlaid on the mesh is a partial sketch of the reduced electrical network. **B**. Example of adjacent partitions re-meshing for equivalent conductance calculation. Partition centroids (orange circles), connecting line (solid black line), and “cut” lines (dashed black lines) are shown. **C**. Solution to Laplace’s equation (Eqn. 1) on the shared volume. **D**. Resulting reduced electrical network, for 200 nodes and the corresponding connections between adjacent partitions. Connection thickness is proportional to the conductance value, and edges in plicate or interplicate regions are shown as orange and blue, respectively. **E**. Solution to Laplace’s equation on the full cleft FEM mesh. **F**. Comparison between voltages on the full cleft FEM mesh and reduced network results. **G**. Schematic of the electrical circuit (Eqn. S2) and illustration of the corresponding electrical network at the ID, equivalent to the network shown in panel D.

Finally, the equivalent conductance between cleft compartments *j* and *k* (*g_c,i,j,k_*), for the cleft between cells *i* and *i* + 1, is calculated by dividing the total current through the mesh, by the voltage difference across the mesh, i.e., *g_c,i,j,k_* = *I_fem_*/*V_fem_*. See Supporting Methods that illustrate this equivalent conductance calculation in a regular rectangular geometry, resulting in the established relationship between electrical conductance, geometry, and electrical resistivity (Fig S4). We repeat this calculation over all pairs of adjacent compartment in the intracellular cleft to obtain the intracleft conductances *g_c,i,j,k_* for all combinations of *j,k* = 1,…,*M* and *j* ≠ *k*. By definition, *g_c,i,j,k_* = 0 for non-adjacent compartments and *g_c,i,j,j_* = 0. Additionally, for all compartments on the ID “periphery,” we calculate the effective conductance between cleft compartment and the bulk extracellular space (*g_b,i,j_*) using a similar procedure, replacing the second compartment centroid with the center of the compartment-ID boundary; for all compartments not adjacent to the bulk, *g_b,i,j_* = 0.

We investigate the dependence of the cleft conductance measurements on intermembrane separation by repeating the mesh generation and cleft conductance measurements for different values of *d_P_* and *d_I_p__*. Additionally, we perform this process five times to replicate the experimental variability in ID and cleft properties. We also investigate the influence of ID structures by measuring cleft conductances in the absence of either GJs or plicate folds.

The resulting electrical network is formed by nodes representing the average electrical potential of the compartment, and edges representing intra-cleft electrical conductance (Fig 3D). We validate this approach by comparing the electrical potential on the reduced cleft electrical network and the corresponding full 3D cleft finite element mesh (calculated in finite element solver COMSOL). We note that the reduced cleft network represents a significant computational reduction: solving for the 3D cleft potential requires solving Laplace’s equation on the entire cleft finite element mesh, comprised of approximately 300,000 finite element tetrahedra (Fig 3E), approximately 1 minute of computational time, while solving for the cleft potential on the reduce cleft network requires solving an *M*-dimensional linear system, obtained from applying Kirchoff’s current law to the network circuit, requiring < 1 second of computational time. Figure 3F illustrates the close comparison of the voltage of the cleft network nodes and the corresponding average partition voltage, for all partitions. In Supporting Material, we quantify the root-mean square error between the cleft network and cleft finite element solution, and we found that increasing the number of partitions *M* reduced the error (Fig S5). Importantly, for 50 or more partitions, we find that the error for the reduced cleft network potential is less than 10%.

### Cleft network tissue model with ID nanodomains

We next developed a novel cardiac tissue model formulation that incorporates the reduced cleft network, thus integrating the effects of ID nanoscale structure into a tissue-scale model. Full details of the cardiac tissue model are provided in Supporting Material, including equations, parameters, and numerical methods. In brief, we simulate a 50-cell cable of ventricular myocytes, with membrane patch dynamics governed by the Luo-Rudy 1 model [41]. As in our prior work [32, 33, 34, 35] and work by others [26, 27, 24, 9, 28, 29], we account for non-uniform Na^+^ subcellular localization by spatially discretizing each cell into an axial membrane patches along the lateral membrane and ID membrane patches at site of the cell-cell junctions. In contrast with prior work in which each ID membrane was represented by a single membrane patch, in this study, each ID is discretized into *M* membrane patches (Fig 3G). The extracellular side of each ID membrane patch, representing the intercellular cleft space, is coupled via the reduced cleft electrical network (Fig 3D, G (right)), as described above. That is, the *j*th and *k*th cleft potential between the *i*th and *i*+1th cell (*ϕ_c,i,j_* and *ϕ_c,i,k_*, respectively) are coupled with conductance *g_c,i,j,k_*. This critical modification incorporates the ID nanoscale structure into the macroscale tissue model.

The intracellular electrical potential (*ϕ_m,i_*) in each cell is coupled to the pre- and postjunctional intracellular potentials (*ϕ_r,i_* and *ϕ_l,i_*, referring to “right” and “left” potentials, respectively) with conductance *g_myo_*. In addition to ephaptic coupling via the shared intercellular cleft, pre- and post-junctional intracellular potentials (*ϕ_r,i_* and *ϕ_l,i+2_*, respectively) are coupled by a GJ conductance *g_gap,i_* Note that the finite element mesh generates the spatial location of individual GJ clusters, such that the tissue model can represent these distinct electrical connections between the pre- and post-junctional intracellular spaces. However, all of the GJ resistors are connected in parallel, such that the overall GJ conductance between neighboring cells can be represented by a single resistor, with conductance equal to the sum of the individual GJ cluster conductances (i.e., the macroscopic GJ conductance). We also note that while in general the cleft conductances (*g_c,i,j,k_*) could differ between different clefts, we assume these values are the same throughout the tissue, i.e., *g_c,1,j,k_* = *g_c,2,j,k_* =… = *g_c,N–1,j,k_*. Similarly we assume the same GJ conductance between all adjacent cells, i.e., *g_gap,i_* = *g_gap_*.

We account for the dynamic [Na^+^] in each of the intercellular cleft compartments ([Na^+^]_*c,j*_), where [Na^+^]_*c,j*_ in the *j*th cleft compartment is governed by the Na^+^ current in the *j*th pre- and post-junctional ID membrane patch, and ionic flux between adjacent compartments (and the bulk, for periphery compartments). The ionic flux between cleft compartments is governed by the electrochemical gradient, i.e., the cleft potential and ionic concentration differences between the adjacent compartments.

We perform simulations with the cleft network tissue model using the cleft conductances obtained from the finite element meshes for different intermembrane separation values. We also vary GJ coupling levels, i.e., values of *g_gap_*. For comparison with the cleft network model, we also perform simulations with a single cleft tissue model, the typical approach of prior studies of ephaptic coupling that neglects ID structural heterogeneity [27, 32, 24], and also with the standard monodomain tissue model formulation, which does not account for the ID, intercellular cleft space, nor non-uniform Na^+^ channel distribution (Fig S6). For all the tissue simulations, we pace the leftmost cell (cell 1) at a cycle length of 500 ms. Conduction velocity (CV) was computed by linear regression of the activation times of the intracellular potential of cells 15 to 35, where the activation time is defined as the time when *ϕ_m_* crosses above −60 mV.

## Results

### Intercalated disk structural properties

Representative TEM images illustrate the interplicate and plicate regions of the ID (Fig 1B), and plicate folds, gap junctions, and intermembrane distance in the plicate and interplicate regions (Fig 1E). Summary measurements of the key ID structures used to generate the finite element meshes are shown in Table 1. In particular, we find several key differences between the plicate and interplicate regions. GJ length is longer and GJs comprise a larger percentage of the cell membrane in the interplicate, compared with the plicate. Intermembrane separation is narrower in the interplicate regions, compared with the plicate, consistent with our recent measurements in atria [20]. Our measurements of plicate fold amplitude are comparable with recent studies as well [2, 3]. Additionally, we find that plicate regions tended to be longer than interplicate, although not significantly.

**Table 1.**
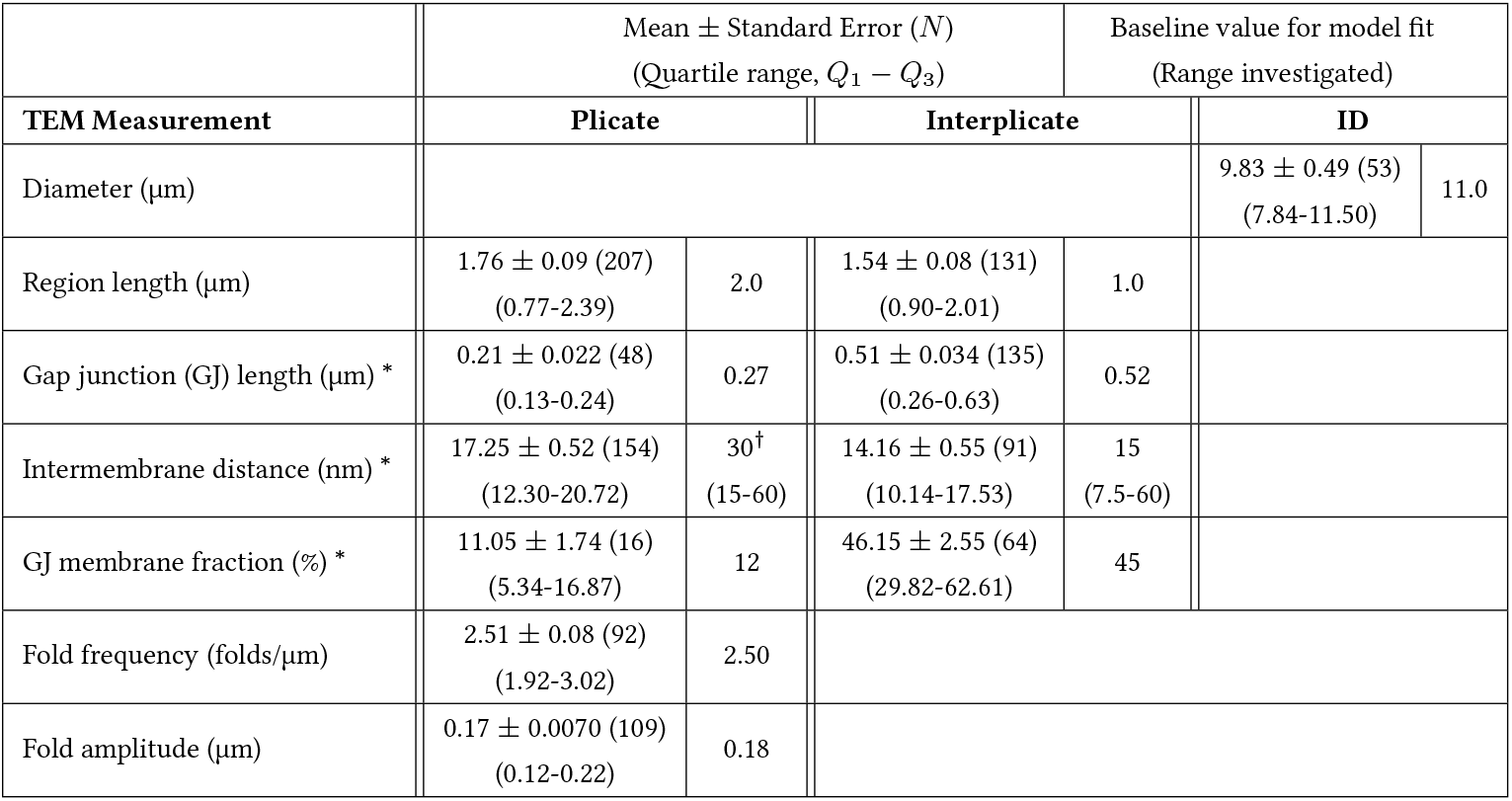
Summary of TEM measurements of ID structures. Data is pooled from images of 3 mouse hearts, and 51 IDs (for intermembrane distance) and 76 IDs (for all other measurements). *N* denotes the total number of measurements. * denotes statistically significant differences (*p* < 0.01) between plicate and interplicate measurements. ^†^ Baseline model value for the plicate intermembrane separation parameter was based on our earlier work in atria [10, 20].

### Intercellular cleft conductances

Using the baseline mesh generation parameters obtained from TEM measurements (Table S1), we first demonstrate the generality of the mesh generation process and created five different ID maps and meshes (Fig. 4A, B). Importantly, each finite element mesh is generated from the same parameters but differ due to randomness inherent in the mesh generation process. Slices through the meshes are visually quite comparable to TEM images, but also differ between each other due to differences in the mesh and orientation of the slices, exhibiting differences in plicate and GJ lengths (Fig 4C). As described above, we calculate the intracleft conductances between all adjacent cleft compartments for each mesh. We divided each mesh into 200 compartments or partitions, resulting in approximately 450 cleft network edges, split between the plicate and interplicate regions. Histograms of the cleft conductances show some small differences between the different meshes (Fig 4D); however, overall the cleft conductance distributions are similar with nearly identical means (black lines).

**Figure 4.**
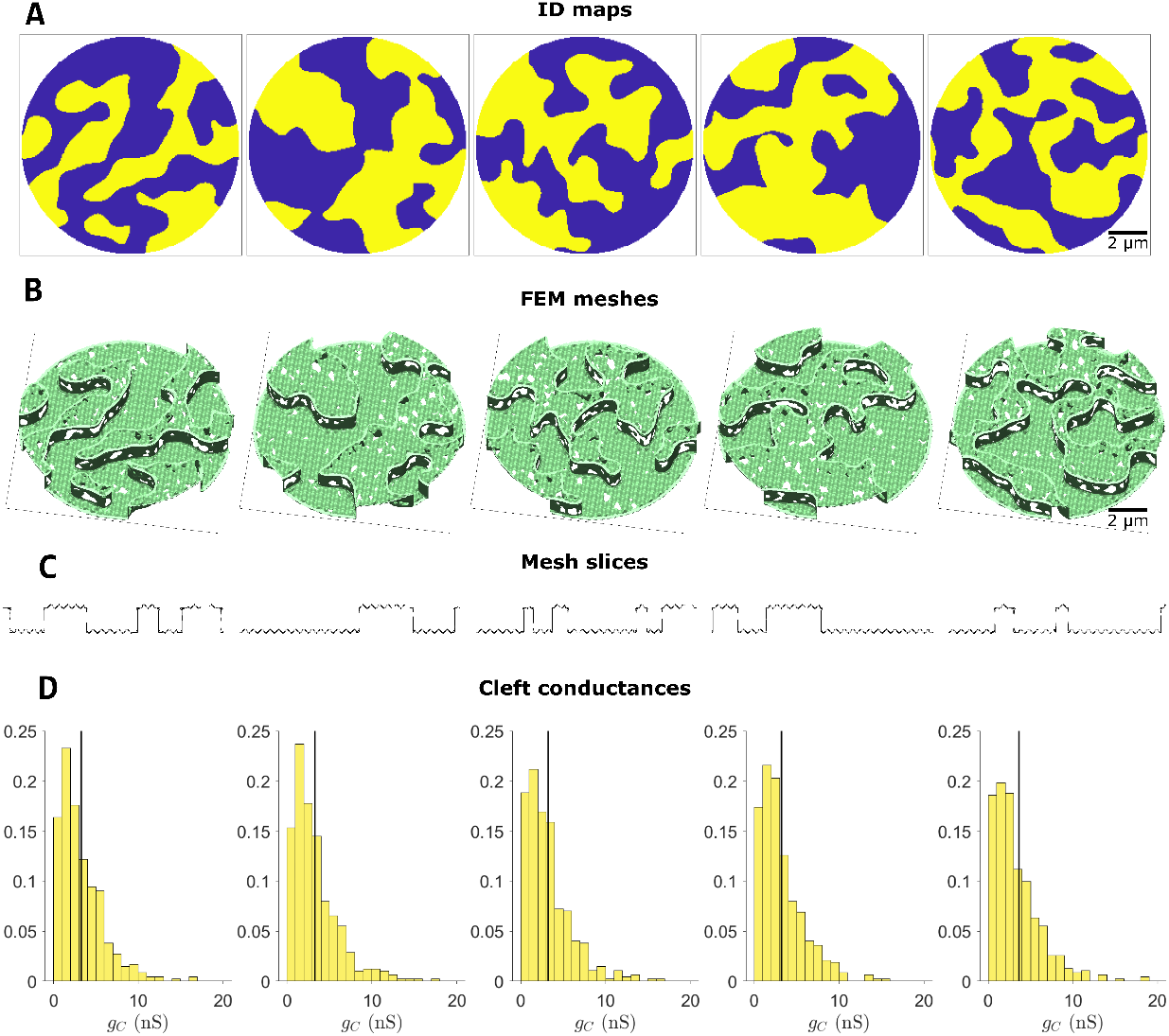
Illustration of multiple cleft finite element model (FEM) meshes and cleft conductance calculations. **A**. Five example intercalated disk (ID) maps, illustrating variable but similar overall geometries. **B**. Corresponding FEM meshes, and **C**. slices through the meshes above, illustrating variation in the lengths of plicate and interplicate regions, and GJs. **D**. Distribution of cleft conductances (*gc*) in each mesh, for M = 200 partitions. The black line represents the mean conductance for each mesh. Parameters: Plicate membrane distance *d_P_* = 30 nm, interplicate membrane distance *d_I_p__* = 15 nm.

We next investigated the differences between cleft conductances specifically within the interplicate (*gI_p_*) and the plicate (*g_P_*) regions, and then further to what extent different ID properties altered these conductances. Histograms illustrate that conductances in the plicate are typically larger than in the interplicate, with the mean plicate conductance (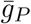, orange vertical line) almost twice as large as the mean interplicate conductance (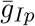, blue vertical line) (Fig 5A, E). Note that these conductances represent all of the values from the five meshes in Figure 4. This overall trend is consistent with higher conductance associated with the wider intermembrane separation in the plicate, compared with the interplicate.

**Figure 5.**
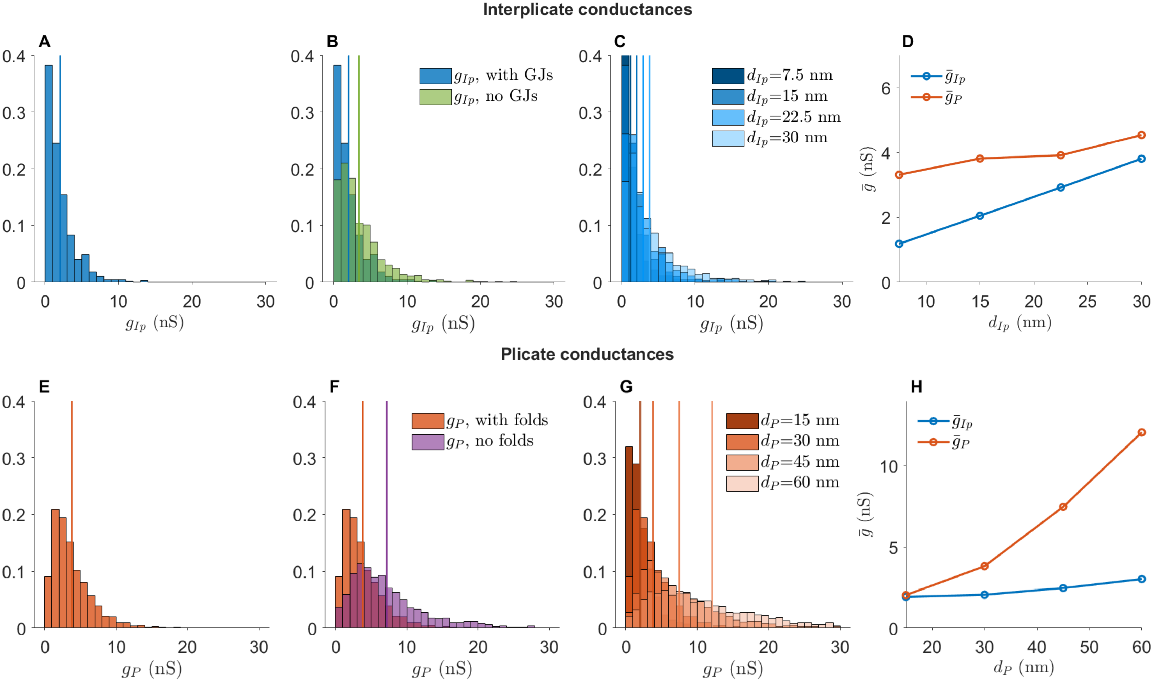
Interplicate and plicate region conductances. **A**. Distribution of conductances in the interplicate region (*gI_p_*), with values from the five meshes in Figure 4. **B**. Interplicate conductances in the presence and absence of gap junction (GJs). **C**. Interplicate conductance distributions for increasing interplicate intermembrane distance *d_I_p__*. **D**. Mean interplicate conductance 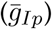 and mean plicate conductance 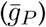 are shown as a function of *d_I_p__*. Note that the blue line values correspond to the vertical lines in C. **E**. Distribution of conductances in the plicate region (*gP*), with values from the five meshes in Figure 4. **F**. Plicate conductances in the presence and absence of plicate folds. **G**. Plicate conductance distributions for increasing plicate intermembrane distance *d_P_*. **H**. 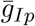 and 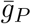 are shown as a function of *d_P_*. Note that the orange line values correspond to the vertical lines in G. In panels A-C and E-G, vertical lines denote distribution means. Parameters: (A-D) *d_P_* = 30 nm, (E-H) *d_I_p__* = 15 nm.

We next investigate to what extent nanoscale ID structures influence cleft conductance, specifically the presence of GJs and plicate folds. We find that GJs decrease conductance in the interplicate (Fig 5B), due their high density in this region. Thus, in addition to providing a direct electrical connection between coupled myocytes, GJs also reduce the electrical conductance of the intercellular cleft. In contrast, as GJs are both smaller and rarer in the plicate, the presence of GJs in the plicate minimally influences cleft conductances (Fig S7D). We also find that plicate folds greatly decrease plicate conductance (Fig 5F), as the folds increase the effective cleft length and thus decrease conductance. As expected, plicate folds do not influence interplicate conductances (Fig S7B).

We additionally investigate how changes in intermembrane separation influence the interplicate and plicate conductances. As expected, increasing interplicate intermembrane separation *d_I_p__* shifts the interplicate conductance histogram to the right (Fig 5C), consistent with an overall increase in mean interplicate conductance 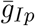 (Fig 5D, blue line). Interestingly, we also observe a small increase in mean plicate conductance 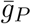 as *d_I_p__* increases, due to a few cleft compartments at the interface between interplicate and plicate (Fig 5D, orange line). Similarly, we find that increasing plicate intermembrane separation *d_P_* shifts the plicate conductance histogram to the right (Fig 5G), consistent with an overall increase in mean plicate conductance 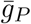 (Fig 5H, orange line), with minimal increase in 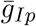 as *d_P_* increases (Fig 5H, blue line).

Overall, the results show that both intermembrane distance and ID structural features alter local intracleft conductance. Interestingly, calculations also predict that the plicate region is more sensitive to changes in intermembrane separation, compared with the inter-plicate. In the absence of irregular structure, conductance would be directly proportional to intermembrane separation. However, simulations predict that a 2-fold increase in *d_P_* (from 30 to 60 nm) leads to a 3.1-fold increase in 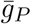, while a similar 2-fold increase in *d_I_p__* (from 15 to 30 nm) leads to a 1.8-fold increase in 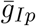. Thus, these changes in sensitivity specifically arise due to structural heterogeneities introduced by GJs and plicate folds. Illustrations of the presence/absence of these ID structures and different intermembrane separation are shown in Figure S8.

### Intercalated disk structure regulation of electrical conduction

We next investigate how nanoscale ID structure and changes in this structure influence tissue level conduction. We first compared our cleft network model with the standard monodomain model and a single cleft tissue model (Fig 6). As expected, the monodomain model illustrates the activation of Na^+^ current during conduction, but does not account for changes in cleft potential and [Na^+^], as these are not accounted for in the formulation (Fig 6, left column). Consistent with prior work [24, 32, 33, 9], the single cleft model illustrates cleft hyperpolarization and a transient depletion of cleft [Na^+^]; additionally, the Na^+^ current magnitude at the ID is increased, relative to the monodomain model, due to the preferential localization of Na^+^ current on the ID membrane (Fig 6, middle column). In contrast with the single cleft model, the cleft network model illustrates the heterogeneous ID Na^+^ currents, cleft [Na^+^], and cleft potentials during conduction (Figure 6, right column).

Importantly, we note that the cleft network average (thick lines) differs from the single cleft model, demonstrating that the single cleft does not capture the overall dynamics of ID currents, cleft concentrations, and cleft potentials. Specifically, we find that cleft potentials are hyperpolarized to a greater extent and cleft [Na^+^] refills slower, compared with the single cleft model, due to the complicated ID geometry and weaker coupling to the bulk. Both of these changes ultimately reduce Na^+^ current driving force, such that Na^+^ current at the ID is also reduced, compared with the single cleft model. In Figure 6E, we illustrate the heterogeneity of cleft polarization during conduction. We find that cleft nodes closer to the ID center are more hyperpolarized than those toward the periphery, due to the low conductance path from the center to the bulk. Further, nodes in the interplicate also tend to be more hyperpolarized than in the plicate, due to lower interplicate conductance (cf. Figure 5A and E).

**Figure 6.**
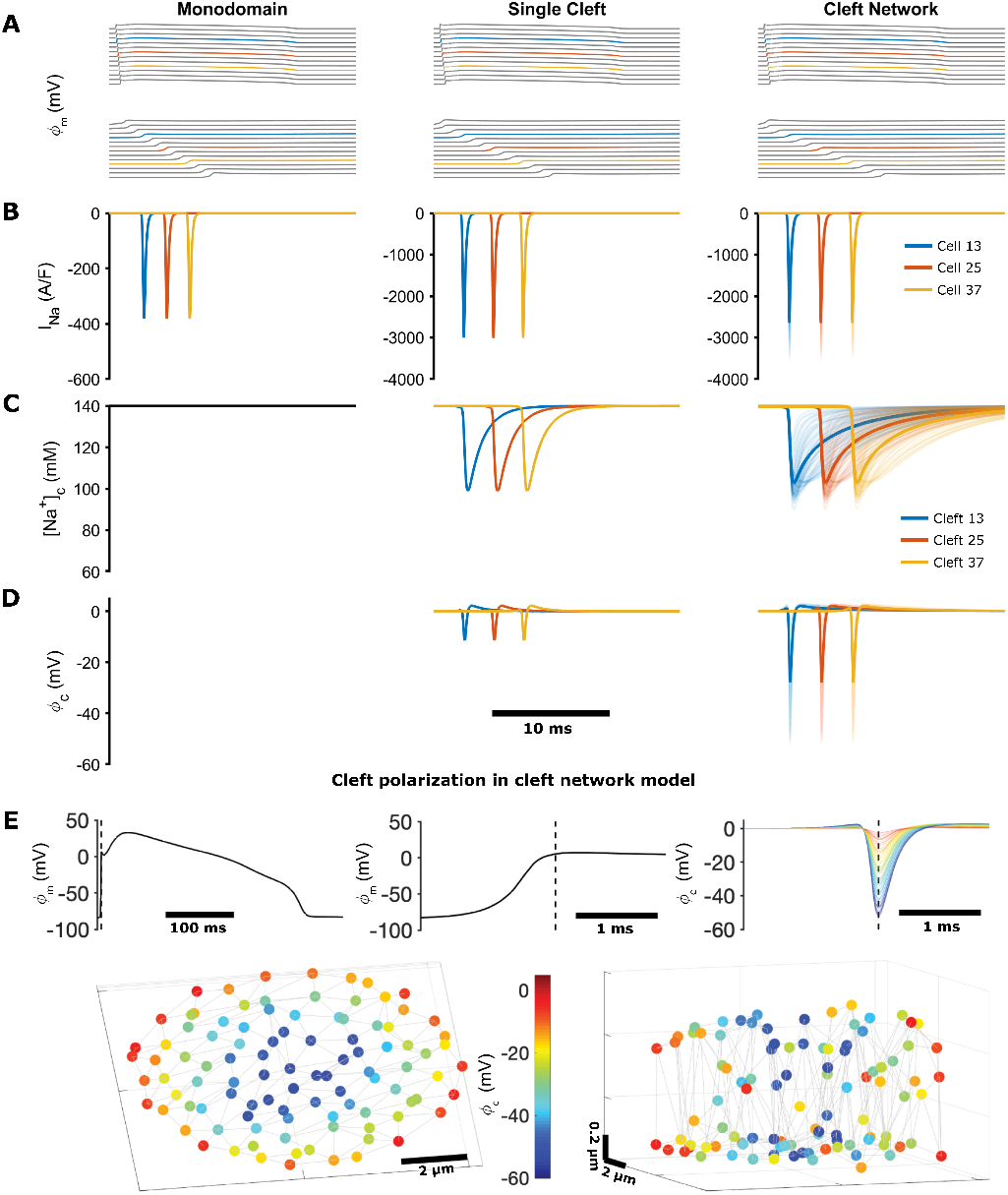
Simulation of conduction in cardiac tissue models, comparing the monodomain model, the single cleft model, and the FEM-derived cleft network model. **A**. Axial membrane transmembrane potential (*ϕ_m_*), illustrating (top) a propagating electrical wave, and (bottom) a magnified view of the action potential upstroke. **B**. Na^+^ current (*I_Na_*) on the axial membrane (monodomain model) or at the ID (single and network cleft models). **C**. [Na^+^] in the intercellular cleft. The monodomain model does not account for dynamic [Na]^+^ or voltage in the cleft. [Na^+^] in all cleft compartments are shown for the cleft network model, with the average shown by the thicker line. **D**. Cleft potential (*ϕ_c_*) in the single cleft and network cleft models, with all compartment potentials and average (thick lines) shown for the cleft network model. In A-D, colored lines represent cell or cleft 13 (blue), 25 (orange) and 37 (yellow), in the 50-cell tissue. **E**. Cleft voltage hyperpolarization in the cleft network model. (Top) Axial transmembrane potential *ϕ_m_* and cleft potential *ϕ_c_* for cell or cleft 25 are shown, with the dashed line indicating the time of maximum cleft hyperpolarization. (Bottom) Color indicates the cleft hyperpolarization at each node in the cleft, with node position indicating the location in the 3D cleft space (shown from two views). Parameters: Single cleft: *d* =15 nm, Cleft network: *M* = 100, *d_P_* = 30 nm, *d_I_p__* = 15 nm.

We next investigate to what extent ID structural differences alter conduction velocity (Fig 7). We measured CV in the cleft network models and compare measurements with the standard monodomain tissue model and a tissue model with a single cleft compartment. Additionally, in the two cleft models, we varied the intermembrane separation for the single cleft or specifically in the interplicate regions for the cleft network (i.e., varying *d_I_p__*). We varied interplicate intermembrane separation, as opposed to varying intermembrane separation throughout the ID, to investigate the influence of perturbations that target specific regions of the ID. We note that the same ID and GJ maps were used to generate finite element meshes with varying interplicate intermembrane separation. Experimental measurements of macroscopic GJ conductance vary considerably, typically 10s to 100s of nanosiemens (nS) [42, 43, 44, 45, 46, 47, 48, 49]. Thus, our simulations consider a range of GJ conductances.

We find that for strong GJ coupling, CV was highest in the monodomain model and lowest in the cleft network model (Fig 7A). By neglecting the ID and non-uniform channel distribution, the monodomain model overestimates CV by approximately 20-30%. The single cleft model also overestimates CV by approximately 10% by neglecting ID structural heterogeneity. For both cleft models, increased intermembrane separation increases CV, to a slightly greater extent in the cleft network model.

For moderate GJ coupling, CV is similar for all three models, although the monodomain model also predicts a slightly faster CV compared with the cleft models (Fig 7B). Interestingly, the cleft models predict different trends for increasing intermembrane separation: CV decreases in the single cleft model and increases in the cleft network model, although these differences are small. Finally, for weak GJ coupling, we find both quantitative and qualitative differences between model predictions (Fig 7C). The single cleft model predicts CV slower than the monodomain model, and CV increasing as intermembrane membrane separation increases. In contrast, CV in the cleft network model is approximately 20% faster, compared with the monodomain model, and CV decreasing as interplicate membrane separation increases. Further, the cleft network model is much more sensitive to changes in the intermembrane separation, compared with the single cleft model, as *d_I_P__* increasing from 15 to 60 nm results in an approximately 20% decrease in CV.

In Figure 7D, we plot CV as a function of GJ conductance (*g_gap_*) for different values of interplicate intermembrane separation (*d_I_p__*). This plot in particular highlights that CV is most sensitive to GJ conductance for wide interplicate intermembrane separation, while in contrast, CV is least sensitive to GJ conductance for narrow interplicate intermembrane separation. Additionally, we observe that CV is most sensitive to interplicate intermembrane separation for either low or high GJ conductance, with opposite trends as noted above. Further, the cleft network model CV, normalized relative to the CV in the monodomain model, is most sensitive to interplicate intermembrane separation for low GJ conductance, demonstrating the regime of the greatest ID structural sensitivity and discrepancy with the monodomain model.

Collectively, these findings show that CV in the cleft network model exhibits a weaker dependence on GJ coupling, in particular for narrow interplicate membrane separation, compared with standard modeling approaches that neglect the ID or assume a homogeneous ID. Further, we find that ID structural heterogeneity can result in enhanced sensitivity to intermembrane separation, compared with the single cleft modeling approach assuming a homogeneous ID and intercellular cleft. We next investigate the mechanism underlying intermembrane separation sensitivity due to ID structural heterogeneity in the cleft network model.

**Figure 7.**
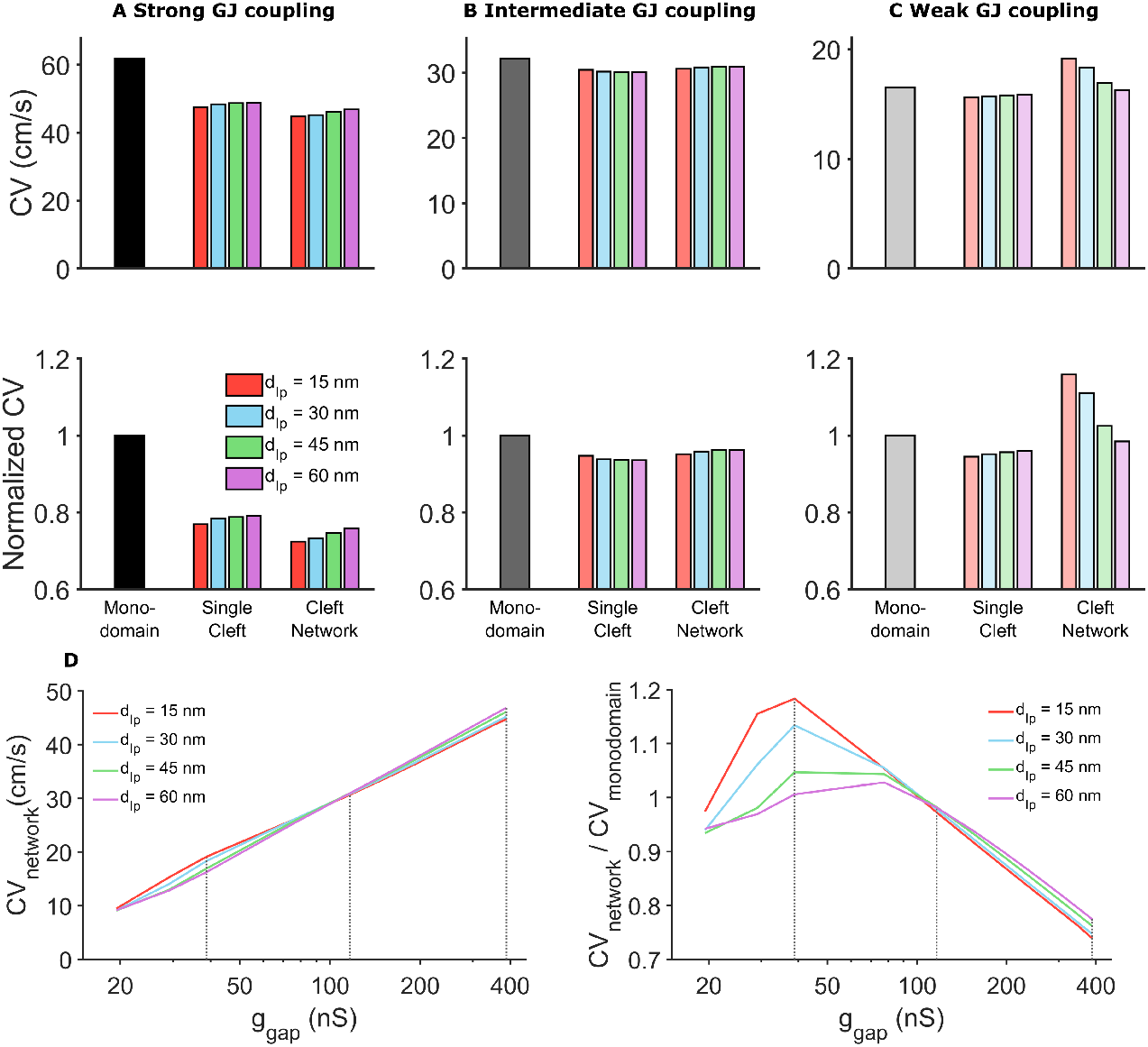
Comparison of conduction velocity (CV) for different tissue models, intermembrane separation, and gap junctional (GJ) coupling strength. **A**. Strong GJ coupling: CV in the single cleft and network models is slower, compared with the monodomain model. In the cleft models, CV slightly increased for increased intermembrane separation (in the single cleft model) and interplicate distance *d_I_p__* (in the cleft network model). **B**. Intermediate GJ coupling: CV is slightly reduced in the cleft models, with minimal dependence on intermembrane separation. **C**. Weak GJ coupling: CV is reduced in the single cleft model but increased in the cleft network model, with different dependence on intermembrane separation. **D**. (Left) For all *d_I_p__* values, CV decreases as gap junctional conductance *g_gap_* decreases. CV is more sensitive to *d_I_p__* for either high or low *g_gap_*, and CV is less sensitive to *g_gap_* for low *d_I_p__*. (Right) Cleft network model CV, normalized to the monodomain CV, is typically greater than 1 for low *g_gap_* and less than 1 for high *g_gap_*. This ratio is highly sensitive to *d_I_p__* for low *g_gap_*. Dotted grey lines corresponding to *g_gap_* values in panels A-C. Note the logarithmic scale for the x-axis. Parameters: *g_gap_* (nS): (A) 388; (B) 116, (C) 38.8. Cleft network, plicate distance *d_P_* = 30 nm. Single cleft, intermembrane separation *d* = *d_I_p__*.

To investigate this sensitivity and the differing responses to interplicate expansion, we consider cleft and membrane dynamics during conduction for four cases (Fig 8): (A) strong GJ coupling and narrow interplicate, (B) weak GJ coupling and narrow interplicate, (C) strong GJ coupling and expanded interplicate, and (D) weak GJ coupling and expanded interplicate. In all cases, we illustrate the intracellular (black) and cleft potentials (blue), cleft [Na^+^] (red), GJ current (orange), and the Na^+^ currents at the pre-junctional (cyan) and post-junctional (magenta) membranes, in the middle of the tissue (between cells 25 and 26).

For strong GJ coupling and narrow interplicate (Fig 8A), the cleft is hyperpolarized in a highly heterogeneous manner (blue, see also Figure 6E), and cleft [Na^+^] (red) is locally depleted, reducing Na^+^ current driving force heterogeneously. Collectively, these result in desynchronized activation of the post-junctional Na^+^ currents, with variable current magnitude (magenta), which reduces the overall ID Na^+^ current density (thick magenta). In contrast, for wider interplicate (Fig 8C), cleft hyperpolarization is reduced and cleft depletion is attenuated, such that post-junctional Na^+^ current is more synchronized and less variable, resulting in a larger ID Na^+^ current density. Additionally, GJ current magnitude (orange) is greater for the wider interplicate. Collectively, the larger Na^+^ and GJ current result in a faster upstroke and thus faster CV for the wider interplicate.

For weak GJ coupling (Fig 8B, D), in both cases, GJ current is greatly reduced and conduction is slower, compared with strong GJ coupling, as expected. Interestingly, expansion of the interplicate exhibits a similar behavior as in the previous case, but with opposite effect on conduction: For narrow interplicate (Fig 8B), heterogeneous cleft hyperpolarization and cleft [Na^+^] depletion similarly result in desynchronized post-junctional Na^+^ current. However, the larger magnitude of cleft polarization results in both an overall earlier activation and longer duration of post-junctional Na^+^ current, compared with a wider interplicate (Fig 8D), which results in an earlier upstroke and faster conduction for the narrower interplicate. Thus, we find that ID structural heterogeneity plays a significant role in regulating conduction: narrow intermembrane separation promotes desynchronized and varying magnitude post-junctional Na^+^ current, which slows conduction for strong GJ coupling yet enhances conduction for weak GJ coupling.

**Figure 8.**
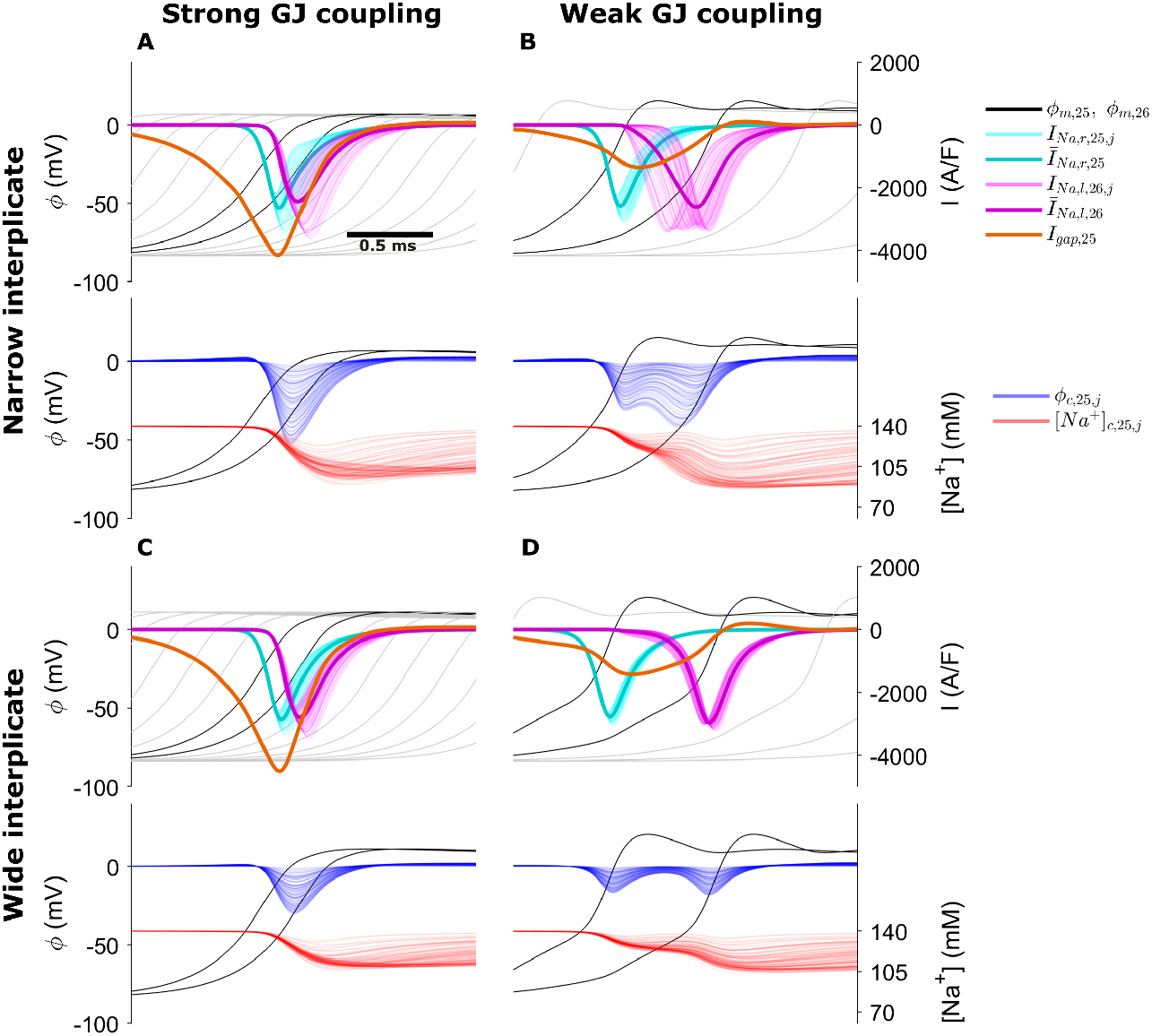
Membrane and cleft dynamics during electrical conduction in the cleft network model for the following cases: **A**. High gap junctional (GJ) coupling, narrow interplicate; **B**. Weak GJ coupling, narrow interplicate; **C**. Strong GJ coupling, wide interplicate; **D**. Weak GJ coupling, wide interplicate. In each panel, (top) the transmembrane potential *ϕ_m_* (black, for cells 25 and 26; gray for upstream and downstream cells), cell 25 pre-junctional (“right,” *I_Na,r_*, cyan) and cell 26 post-junctional (“left,” *I_Na,l_*, magenta) Na^+^ current, GJ current (orange); (bottom) cleft potential *ϕ_c_*, and cleft [Na^+^] for all cleft compartments are shown as a function of time. Dark cyan and magenta curves represent the mean pre- and post-junctional *I_Na_*, respectively. Parameters: *M* = 100, Plicate distance *d_P_* = 30 nm, Gap junctional conductance *g_gap_* (nS): (A, C) 388; (B, D) 38.8, interplicate distance *d_I_p__* (nm): (A, B) 15, (C, D) 60.

## Discussion

In this study, we developed an approach to generate realistic FEM meshes of the ID, based on TEM-derived measurements of nano- to microscale structure, which are incorporated into a 1D tissue model of cardiac conduction. To our knowledge, this is the first study to integrate FEM-based nanoscale structural modeling with a tissue-scale cardiac electrophysiology model. We investigated the effects of ID structure on intracleft conductance on multiple FEM mesh samples: we found that conductance increases with intermembrane distance, with differences between distinct regions (i.e., plicate vs. interplicate); and that conductance in interplicate regions is decreased by the nanoscale structure of gap junctions and associated perinexi [16, 50, 9], while membrane folds have a similar effect in plicate regions.

By incorporating FEM mesh structural data into a tissue model, we demonstrate that changes in regional ID intermembrane distance (i.e. in the interplicate regions) regulate cardiac conduction. In particular, we found that for tissue with strong GJ coupling, interplicate region intermembrane expansion increased CV, while for tissue with weak GJ coupling, we observe the opposite relationship. Importantly, we found that both effects depend on the ID and cleft structural heterogeneity: specifically, heterogeneous cleft hyperpolarization leads to desynchronized activation of the post-junctional Na^+^ currents. This heterogeneous behavior, for strong GJ coupling, leads to a lower average Na^+^ current and slower CV. However, in the case of weak GJ coupling, this behavior activates the downstream cell earlier and thus increases CV. More broadly, we find that previous model formulations that neglect ID structural heterogeneity (i.e., the single cleft model) or the ID entirely (i.e., the monodomain model) can either under- or overestimate predictions of CV. Overall, the cleft network model exhibits a weaker dependence of CV on GJ coupling, compared with other modeling approaches. In particular, across a wide range of GJ coupling strengths, we find that CV is less sensitivity to GJ coupling for narrower interplicate intermembrane separation, while in contrast CV sensitivity to GJ coupling is enhanced for wider interplicate intermembrane separation.

Our model predictions demonstrate two critical points with implications for predicting arrhythmia risk in physiological and pathological settings, specifically i) that incorporating ID structural details into cardiac tissue models can impact conduction predictions (compared with prior approaches), and ii) that perturbations in ID structure (e.g., inter-plicate intermembrane expansion) can significantly alter conduction. This second point is consistent with prior experimental studies demonstrating that ID disruption can alter conduction and arrhythmia risk. For example, recent work has shown perinexal structure (interplicate ID) is perturbed in atrial tissue from patients atrial fibrillation [23]. We also recently showed that vascular leak, induced by inflammatory cytokine VEGF, disrupts ID nanodomains in both GJ- and MJ-adjacent regions, and ultimately slows conduction and increases susceptibility to arrhythmias [20]. We previously showed that disruption of GJ-adjacent perinexi via edema [19, 9, 51, 21], inflammation [21], or adhesion-disrupting peptides targeting the Na^+^ channel β1 subunit [10] can also slow conduction. In agreement with the majority of GJs residing in the interplicate regions, these conduction slowing trends are consistent with our findings in the setting of weaker GJ coupling and interplicate expansion (Fig. 7C). We speculate that differences between experiments and simulations for cases of stronger GJ coupling may arise due to the lack of accounting for intra-cleft clustering of Na^+^ channels at GJ-adjacent perinexi [10,9,11] and MJ-associated nanodomains [5, 20] within the different ID regions [16, 9]. Incorporating these details is a focus of future work.

We further note that GJ coupling decreases in the diseased myocardium, in cases of heart failure [52, 53] or myocardial ischemia [54]. Our simulation results, particularly for the weak GJ coupling case, may help resolve apparent discrepancies between the time course of GJ remodeling and conduction slowing in the failing heart [52, 53], where edema is known to play a role [55]. We therefore expect that CV sensitivity to interplicate intermembrane separation (shown in Fig. 7) is important in modeling these pathologies, especially in a heterogeneously affected tissue. One important direct extension of our current work is to investigate mechanisms of electrical dysfunction in failing hearts. Importantly, our approach is sufficiently flexible to incorporate ID structural data from failing hearts into FEM and tissue models, and such mechanistic studies is a key focus of future work.

There are a number of prior studies that quantify ID structure. Our measurements of overall ID size [56, 57], intermembrane spacing and GJ cluster size [58] and plicate folds and amplitude [3] are quite similar to previously published data. In contrast with our approach, other studies have characterized the entire 3D structure of the ID using 3D stacking and image segmentation [1, 2, 3]. We note that our approach of mesh partitioning and calculating equivalent cleft conductances, which could be then incorporated into the cleft network tissue model, can be applied to a 3D FEM model based on a 3D ID reconstruction. However, our approach is more flexible and has a number of specific advantages over a direct 3D reconstruction. First, we are able to directly modify specific features of the ID mesh geometry, to investigate the role of such features on conduction. A comparable 3D reconstruction would be required for each experimental case, and as 3D stacking and segmenting images is quite time-consuming and challenging, this would greatly limit the conditions investigated. Second, our approach can determine ID geometry based on hundreds of images from multiple hearts and IDs, thus accounting for physiological variability, while a 3D reconstruction only provides information on the specific ID being imaged. Finally, our approach is not limited to TEM images, as we could derive FEM parameters from any imaging modality, provided that there is sufficient resolution to characterize the relevant ID structures.

In a prior modeling study, Hichri and colleagues simulated two equally-spaced apposing cell membranes, separated by an intercellular cleft space modeled using a uniform FEM mesh [37]. They similarly predict that cleft potential (*ϕ_c_*) is variable within the cleft, specifically with larger magnitude hyperpolarization towards the center (a result also described by Mori et al [25] in a 3D radially-symmetric model). This variability in turn drives a heterogeneous activation of the postjunctional membrane, similar to our results in Figure 8, and the authors show that these effects are amplified by both decreased intermembrane distance and Na^+^ channel clustering. We similarly find that the cleft is more polarized towards the center (Fig 6). We further predict that the potential distribution is in fact even more variable and heterogeneous due to the irregular structure of the ID meshes and variations in structure that arise due to differences in intermembrane separation in the plicate and interplicate and the presence of GJs and plicate folds. Additionally, our approach to derive an equivalent reduced electrical circuit to represent the cleft conductances eliminates the computationally costly step of simulating electrochemical dynamics on the full 3D FEM mesh, facilitating simulation of cardiac tissue and predictions of CV.

Interestingly, the mechanism of conduction slowing/enhancing shown in Figure 8 is similar to the single cleft model previously predicted by Kucera and colleagues [27]: For strong GJ coupling, narrow intercellular clefts enhance cleft polarization and thereby reduce Na^+^ current driving force, which slows conduction, a mechanism described as “self-attenuation.” In contrast, for weak GJ coupling, narrow clefts activate the post-junctional membrane earlier and enhance conduction. Our study builds on this prior work and predicts that cleft heterogeneity also contributes to these mechanisms. Specifically, for strong GJ coupling, self-attenuation and reduced Na^+^ current also arise due to desynchronized activation of the post-junctional membrane. For weak GJ coupling, narrow clefts enhance cleft polarization heterogeneity that facilitates an overall earlier activation of post-junctional Na^+^ channels. Additionally, this earlier activation in one region of the post-junctional ID membrane in turn contributes to the activation of additional Na^+^ channels in other regions, similar to the mechanism of “self-activation” described by Hichri et al [37]. In addition to illustrating regulation by ID structural heterogeneity, our study illustrates that these mechanisms of conduction regulation can be mediated by intermembrane separation changes in a specific region of the ID, specifically the interplicate. Our on-going work is investigating how additional ID structural changes impact conduction.

We conclude by acknowledging limitations of our study. Although our model formulation incorporates significant subcellular details in the representation of cardiac tissue, this representation is still a simplification of the complex cardiac tissue structure. Specifically, our tissue model represents a 1D strand of coupled cells and thus does not account for the 3D tissue structure of the heart. However, importantly our model formulation does represent key subcellular and structural details, specifically non-uniform Na^+^ channel distribution and ID structure. Further, our approach can modified to account for specific ID structural heterogeneity that is not specifically represented in TEM images. One such example are “ribbon” GJs, large GJs that have been identified towards the periphery of the ID [59, 60]. Although we did not measure these directly in our TEM images, we can investigate their potential role in modulating conduction. To illustrate this approach, we created a FEM mesh with a distribution of ribbon GJs on the ID periphery (Fig. S9), and interestingly found that these larger GJs reduced connection to the extracellular bulk, and thus amplify the CV sensitivity to changes in intermembrane distance. More broadly, our work can be further extended to incorporate structural details from other imaging modalities, e.g., ion channel clustering within GJ- and MJ-associated nanodomains, as noted above.

Further studies are also needed to investigate the role of additional electrogenic proteins that have been identified to be preferentially localized at the ID, in particular K^+^ channels [17, 61, 51]; here we focused on the role of Na^+^ channels and their role in regulating conduction at the ID. Finally, while accounting for the irregular structure of the ID membrane, we assume that the lateral membrane follows an overall cylindrical shape, and prior work has shown that microscale heterogeneity, in particular in cell shape/geometry and GJ protein distribution, can impact conduction delays and lead to heterogeneous conduction [62, 63, 64]. We are interested in how these nano- and microscale heterogeneities can influence each other and impact conduction, in particular in pathological settings and associated structural remodeling. Overall, while these limitations may account for quantitative differences in model predictions, importantly our study demonstrate a novel mechanism in which ID nanoscale structural heterogeneity modulates ID-localized Na^+^ channels and regulates cardiac conduction.

## Acknowledgements

This study was supported by funding from the National Institutes of Health, grant numbers R01HL138003 (SHW) and R01HL148736 (RV).

